# The cold tolerance of an adult winter-active stonefly: How *Allocapnia pygmaea* (Plecoptera: Capniidae) avoids freezing in Nova Scotian winters

**DOI:** 10.1101/2025.08.05.668668

**Authors:** Jona Lopez Pedersen, Luke S. Burton, Tamara M. Rodela, Jantina Toxopeus

## Abstract

*Allocapnia pygmaea* Burmeister (Plecoptera: Capniidae) is a winter-active stonefly in North America. Despite the adult’s winter emergence, little is documented about the cold tolerance and cryoprotective biochemistry of *A. pygmaea*. To better understand the cold tolerance of this winter-active stonefly, we collected adult *A. pygmaea* in Antigonish, NS during March or April in both 2023 and 2024. Following different cold exposures, we measured the lower limits of activity (–8.9 °C) and the temperature at which internal freezing occurs (–11.9 °C), and confirmed that *A. pygmaea* could survive subzero temperatures unless they froze. We also assayed the following putative cryoprotectants: proline, glycerol, *myo*-inositol, trehalose, and glucose in control (field-collected) and cold-shocked groups. We detected little effect of cold shock on most cryoprotectants, except for the polyols glycerol and *myo*-inositol, which decreased in concentration following cold shock. These findings improve our current understanding of Capniid cold tolerance, confirming that *A. pygmaea* uses a freeze-avoidant strategy, and lay a foundation for future studies on how they may use cryoprotectants for winter-activity.

## Introduction

*Allocapnia pygmaea* (Burmeister) is a winter stonefly (Plecoptera: Capniidae) distributed in North America, notable for its winter-active adult stage (Hilsenhoff 2001). Little is documented about its life as it transitions from an aquatic nymph to a terrestrial adult in winter and early spring, despite an ability to crawl through ice crevices and walk on top of snow (Moore and Lee 1991). While the effect of low temperature on survival has been partially characterized in adult *A. pygmaea* in Minnesota, USA (Bouchard et al. 2009), the limits to their activity at low temperatures and the mechanisms underlying their cold tolerance have not been studied.

Low temperatures (e.g., near or below 0°C) impair the activity and survival of insects in multiple ways. As environmental temperature decreases, insects will first lose neuromuscular function (ability to move) at a species-specific critical thermal minimum (CT_min_) temperature (Overgaard and MacMillan 2017). Their internal fluids will then freeze at the supercooling point (SCP) temperature (Sinclair et al. 2015), which may be equal to (e.g., Schoville et al. 2015; Toxopeus et al. 2016) or below (e.g., Clarke et al. 2013) the CT _min_. While freeze-tolerant insects survive internal freezing (e.g., Ramløv et al. 1992), freeze-avoidant species do not (e.g., Golding et al. 2023; Schoville et al. 2015). The lowest temperature an insect can survive (i.e., its lower lethal temperature; LLT) is equal to the SCP for freeze-avoidant insects, below the SCP for freeze-tolerant insects, and above the SCP for chill-susceptible insects (Sinclair et al. 2015).

Low molecular weight cryoprotectants are important for both freeze avoidance and freeze tolerance, and can fulfill different functions (Overgaard and MacMillan 2017; Toxopeus and Sinclair 2018). Insects may accumulate cryoprotectants either prior to or in response to cold stress. Increased concentrations of cryoprotectants depress the insect’s SCP, which is a key mechanism for freeze-avoidant insects to prevent internal ice formation (Feng et al. 2016; Toxopeus and Sinclair 2018). Cryoprotectants may also directly protect cells and their macromolecules from challenges associated with low temperatures and freezing (Teets and Denlinger 2013; Toxopeus et al. 2019). During its nymph stage, a freeze-tolerant stonefly, *Nemoura arctica* Esben-Petersen (Plecoptera: Nemouridae), continuously produces glycerol while frozen (Walters et al. 2009). Many cryoprotectants (trehalose, glucose, proline, glycerol) are also metabolites that can be broken down to facilitate ATP production (Jørgensen et al. 2023; Thompson et al. 2003; Toxopeus et al. 2019). To our knowledge, the cryoprotective biochemistry of *A. pygmaea* has not previously been described.

In this study, we characterize the activity and survival of field-collected *A. pygmaea* at subzero temperatures and report concentrations of common insect cryoprotectants, improving our understanding of how an adult winter-emerging stonefly survives Nova Scotian winters.

## Methods

We collected adult *A. pygmaea* from Antigonish, NS, Canada (45.62 °N, – 62.00 °W) in the spring (March to April) of 2023 and 2024 from the walls of a residence near Brierly Brook, a mid-sized stream. Using a paintbrush, we gently transferred insects into 50 mL Falcon tubes containing moist paper towel, transported them inside a cooler with ice packs to St. Francis Xavier University, and housed them in a 4 °C fridge within an hour of collection. *A. pygmaea* destined for metabolite experiments were flash-frozen and stored in a –80 °C freezer for future cryoprotectant assays, while *A. pygmaea* destined for whole-animal experiments remained in the 4 °C fridge for 3-7 days. We weighed stoneflies with an XPR2 microbalance (Mettler Toledo, Columbus, OH, USA) prior to the manipulations described below.

To determine CT_min_ of *A. pygmaea*, we cooled eight stoneflies (2024 collection) from 4 °C at –0.25 °C/min to –12 °C in individual transparent chambers, and classified stonefly activity every 5 min using the same equipment described for measuring CT_min_ in Adams et al. (2025a). Stoneflies were classified as active if they were clearly motile or maintaining an upright position on the wall of the observation chamber. They were classified as inactive if they had fallen to the bottom of the observation chamber. Two stoneflies never fell to the bottom of the chamber, but remained in the same position on the tube wall for more than 100 minutes (appeared “stuck”), and were therefore also classified as inactive. We defined CT_min_ as the temperature at which each stonefly first became inactive, and remained so for at least three observation periods. We tested whether CT_min_ varied as a function of stonefly mass using a linear regression in R (v4.2.2).

To determine SCP, cold tolerance strategy, and survival following acute cold shock, we exposed stoneflies to low temperatures in individual containers using the same equipment described in Adams et al. (2025b). To measure SCP, 28 stoneflies (2023 collection) were cooled from 4 °C to –20 °C at –0.25 °C/min. We then tested whether SCP varied as a function of stonefly mass using a linear regression in R (v4.2.2). To determine cold tolerance strategy, stoneflies (2023 collection) were cooled from 4 °C at –0.25 °C/min to a temperature (ca. –11.5 °C) at which half of them (7 of 16) had frozen and the remaining stoneflies were unfrozen. We inferred cold tolerance strategy based on survival of these two groups after 24 h at 4 °C (Sinclair et al. 2015). To determine their ability to survive acute cold shocks and further verify cold tolerance strategy, we exposed groups of 7 or 8 stoneflies (2024 collection) for 60 min to one of five temperatures ranging from c. –5 to –11.5 °C (encompassing 0 to 100% mortality), and assessed survival after 24 h at 4 °C. Stoneflies were classified as alive if they could move or respond to a gentle physical stimulus (touching with a paintbrush) when returned to room temperature.

We determined the concentrations of five putative cryoprotectants in whole-body homogenates of *A. pygmaea* (2024 collection) via spectrophotometric assays. Each stonefly was homogenized individually with a plastic pestle in 100 μL of Tris-buffered saline (TBS; 5 mM Tris; 137 mM NaCl; 2.7 mM KCl, pH 6.6). We centrifuged them for 5 min at 3000 × *g* and 4 °C. We centrifuged 100 μL of supernatant for 30 min at 3000 × *g* and 4 °C. Supernatants were aliquoted into 1.7 mL tubes for subsequent cryoprotectant assays. We began proline assays immediately post-centrifugation following Carillo and Gibon (2011), while remaining heat-treated (70 °C for 10 min) aliquots were flash-frozen in liquid nitrogen and stored at –80 °C for up to 3 weeks prior to processing. We used the K-INOSL and K-GCROLGK kits (Megazyme, Bray, Ireland) to assay *myo*-inositol and glycerol, respectively, following the manufacturer’s instructions. For trehalose and glucose, we followed Tennesen et al. (2014), using the porcine trehalase and glucose assay reagent (Sigma Aldrich, Toronto, ON, Canada). For all assays, we performed a two-fold dilution series to produce a standard curve. To read absorbances, we used a SpectraMax iD3 (Molecular Devices, San Jose, CA, USA) spectrophotometer. We compared cryoprotectant concentrations in 16 field-collected stoneflies and 8 that were cold-shocked at –4.9 °C for one hour, using ANCOVA tests with mass as a covariate in R (v4.2.2).

## Results & Discussion

We showed that *A. pygmaea* were active at subzero temperatures, but did not survive freezing, demonstrating freeze avoidance. Interestingly, most (6 of 8) stoneflies remained active until they froze, with a CT_min_ between –8.1 and –10.2 °C (Fig. 1A). This is similar to some other winter-active insects that also retain neuromuscular function until they freeze (e.g., Schoville et al. 2015; Toxopeus et al. 2016). In a separate trial, we found that the SCP of stoneflies ranged from –6.0 to –15.0 °C (Fig. 1A), with a mean (–11.9 °C) similar to that of the same species studied in Minnesota, United States of America (Bouchard et al. 2009). Neither CT_min_ (F_1,6_ = 0.67, P = 0.445) nor SCP (F_1,24_ = 0.18, P = 0.673) correlated with mass. While the Minnesota study suggested *A. pygmaea* may be chill-susceptible (Bouchard et al. 2009), our work supports that they are cold-tolerant and can survive temperatures near their SCP as long as internal freezing does not occur (Table 1). When we gradually cooled stoneflies to a temperature at which half froze and half remained unfrozen, all frozen individuals died, while all, unfrozen individuals survived (Table 1). When stoneflies were exposed to subzero temperatures for longer (1 h), all frozen individuals died and almost all (96%) unfrozen individuals survived, regardless of the specific temperature (Fig. 1B). These cold tolerance parameters suggest *A. pygmaea* are freeze-avoidant and can remain active and survive at temperatures they are likely to experience in their environment. For example, in March 2024, air temperatures in Antigonish ranged between –12.8 °C to 8.4 °C (Environment Canada 2024), and the lowest temperatures could be avoided by moving to warmer microenvironments (closer to 0 °C) under snow.

**Table 1.** Cold tolerance strategy of *Allocapnia pygmaea* from Antigonish, NS, Canada after cooling (2023 collection) or cold-shock (2024 collection).

| Collection<br>year | Mean mass<br>± SE (mg) | *N* survived/<br>*N* frozen | *N* survived/<br>*N* supercooled | Cold tolerance<br>strategy |
| --- | --- | --- | --- | --- |
| 2023<sup>a</sup> | 4.41 ± 0.94 | 0/7 | 8/9 | Freeze-avoidant |
| 2024<sup>b</sup> | 2.92 ± 0.19 | 0/12 | 24/25 | Freeze-avoidant |

*N* = number of stoneflies; SE = standard error; <sup>a</sup>all stoneflies cooled to -11.5 °C; <sup>b</sup>data
summarized from multiple temperatures (see Fig. 1B).

**Figure 1.**
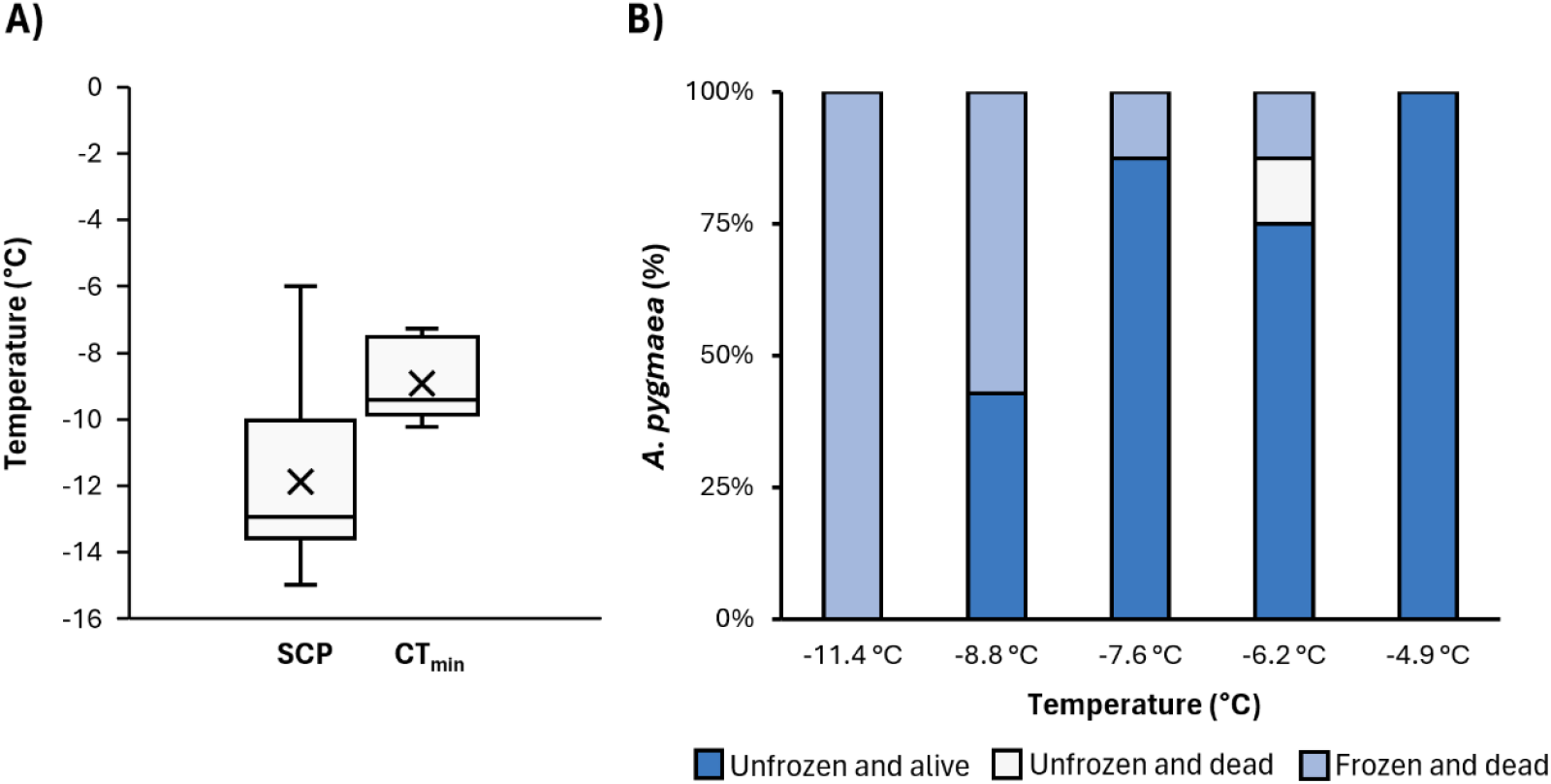
Cold tolerance of *Allocapnia pygmaea* collected in Antigonish, NS, Canada in spring 2023 and 2024. (A) Critical thermal minimum (CT_min_) and supercooling point (SCP). For each interquartile range, the middle line indicates the median while the top represents the first quartile and the bottom represents the third. X indicates the mean. (B) Proportion survival following 1 hour exposure to one of five low temperatures. Each stonefly was categorized based on whether it remained supercooled (unfrozen) or froze during the temperature treatment, and whether it survived or died within 24 h following the treatment.

Other insects that are winter-active or survive in low-temperature environments demonstrate diverse cold tolerance strategies. Like *A. pygmaea*, the insects of the genera *Chionea* Dalman (Diptera: Tipulidae) and *Grylloblatta* Walker (Grylloblattodea: Grylloblattidae) are winter-active and freeze-avoidant (Golding et al. 2023; Schoville et al. 2015). Both inhabit alpine environments with variable temperature ranges but seek out microclimates as low as –3 °C. However, snow flies continue to move at temperatures as low as –10 °C and may even resort to self-amputation of limbs to avoid lethal freezing (Golding et al. 2023). However, some winter-active insects are freeze-tolerant, such as *Hemideina maori* Pictet et Saussure (Orthoptera: Stenopelmatidae) of New Zealand’s mountain habitats (Ramløv et al. 1992; Sinclair et al. 1999) and nymphs of the Arctic stonefly *N. arctica* (Walters et al. 2009). While cold tolerance is not yet investigated in *A. pygmaea*’s nymph stage, it may differ from the adult’s freeze avoidance.

In our assessment of putative cryoprotectants, field-collected stoneflies had the highest concentration of *myo*-inositol, while glycerol, sugars, and proline were less abundant (Fig. 2). The concentration of *myo*-inositol was at least a 3-fold higher (ca. 40 μmol/g) than any other metabolite we measured. *myo*-Inositol is a less commonly reported cryoprotectant in insect cold tolerance literature, with some exceptions (e.g., Štětina et al. 2025; Toxopeus et al. 2019; Vesala et al. 2012). Glycerol (ca. 15 μmol/g) was second most abundant. This is in contrast to freeze-tolerant Arctic stonefly nymphs, whose primary cryoprotectant is glycerol, which accumulates when the insects are frozen (Walters et al. 2009). Other polyols—like threitol, sorbitol, and ribitol—also serve a cryoprotective role in cold-tolerant insects (e.g., Miller and Smith 1975; Hamilton et al. 1985), and may be worth investigating in future analyses of stoneflies. Trehalose, glucose, and proline had the lowest concentrations (< 10 μmol/g) in *A. pygmaea*, which is congruent with work on *N. arctica* nymphs showing that they have detectable but not high concentrations of proline and trehalose (Walters et al. 2009). Cryoprotectant abundance also often declines in spring (e.g., Feng et al. 2016; Vesala et al. 2012), so earlier measurements of cryoprotectants in *A. pygmaea* could show different trends.

**Figure 2.**
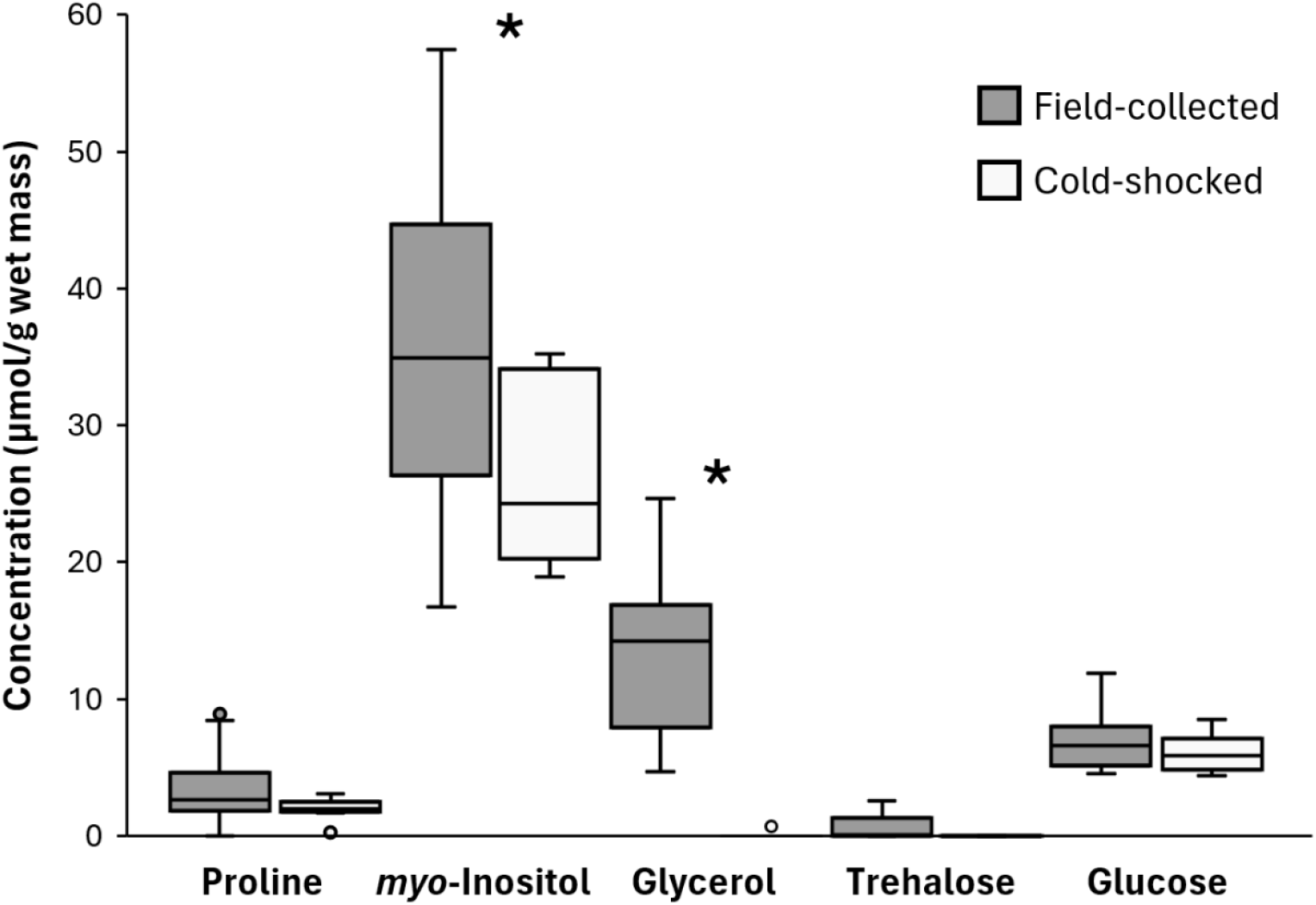
Concentrations (μmol/g) of putative cryoprotectants in whole-body homogenates of *Allocapnia pygmaea* collected in Antigonish, NS, Canada in March 2024. For each interquartile range, the middle line indicates the median while the top represents the first quartile and the bottom represents the third. Outlier points are represented by dots. An asterisk indicates a significant decrease between field-collected and cold-shocked stoneflies, as determined by an ANCOVA test (Table 2).

While short, nonlethal temperature exposures often cause insects to accumulate cryoprotectants (as part of a “rapid cold hardening” response), the stoneflies in our study seemed to deplete putative cryoprotectant reserves following a cold shock. When stoneflies were cold-shocked at –4.9 °C and allowed to recover for 24 h at 4 °C, there was a trend of decreased concentration for all putative metabolites, which was statistically significant for glycerol and *myo*-inositol (Fig. 2, Table 2). As a short-lived, non-feeding adult (Hilsenhoff 2001), it may be adaptive for *A. pygmaea* to use glycerol as a source of ATP (e.g., via pathways described in Thompson et al. 2003; Jørgensen et al. 2023) to recover from injury and/or move towards a warmer microclimate following exposure to subzero temperatures. The exact role of cryoprotectants across life stages is something future studies may tackle, expanding our knowledge on an understudied but bioindicative and climate-sensitive species (Agouridis et al. 2015).

**Table 2.** Effect of treatment (field-collected or cold-shocked) on cryoprotectant concentrations in *Allocapnia pygmaea* collected in Antigonish, NS, Canada in March 2024. Asterisks indicate a significant *P* value from ANCOVA test with mass as a covariate.

| Cryoprotectant | Cold Treatment |  | Mass |  |
| --- | --- | --- | --- | --- |
|  | *F* | *P* | *F* | *P* |
| *myo*-Inositol | F<sub>1,21</sub> = 4.91 | < 0.050 * | F<sub>1,21</sub> = 24.55 | < 0.001 * |
| Glycerol | F<sub>1,21</sub> = 38.15 | < 0.001 * | F<sub>1,21</sub> = 5.26 | < 0.050 * |
| Trehalose | F<sub>1,21</sub> = 3.37 | 0.080 | F<sub>1,21</sub> = 1.22 | 0.281 |
| Glucose | F<sub>1,21</sub> = 0.76 | 0.393 | F<sub>1,21</sub> = 3.97 | 0.060 |
| Proline | F<sub>1,21</sub> = 3.17 | 0.090 | F<sub>1,21</sub> = 3.09 | 0.093 |

## Acknowledgements

The authors would like to thank S.E. Rokosh for assistance with field collections. JLP and LSB were each supported by a Nova Scotia Graduate Scholarship, and the research was supported by a Natural Sciences and Engineering Research Council of Canada (NSERC) Discovery Grant to JT.

